# Systematic classification of shared components of genetic risk for common human diseases

**DOI:** 10.1101/374207

**Authors:** Adrian Cortes, Calliope A. Dendrou, Lars Fugger, Gil McVean

## Abstract

Disease classification is fundamental to clinical practice, but current taxonomies do not necessarily reflect the pathophysiological processes that are common or unique to different disorders, such as those determined by genetic risk factors. Here, we use routine healthcare data from the 500,000 participants in the UK Biobank to map genome-wide associations across 19,628 diagnostic terms. We find that 3,510 independent genetic risk loci affect multiple clinical phenotypes, which we cluster into 629 distinct disease association profiles. We use multiple approaches to link clusters to different underlying biological pathways and show how these clusters define the genetic architecture of common medical conditions, including hypertension and immune-mediated diseases. Finally, we demonstrate how clusters can be utilised to re-define disease relationships and to inform therapeutic strategies.

**One sentence summary:** Systematic classification of genetic risk factors reveals molecular connectivity of human diseases with clinical implications

---

The classification of human diseases is central to their diagnosis, prognosis and treatment but poses a long-standing challenge. Current clinical practice is largely organised by the tissues or organs affected, but this distinction does not necessarily reflect the biological relationships that connect or segregate diseases based on their underlying pathophysiology (*1*). Genome-wide association studies (GWAS) of risk for common medical conditions have revealed widespread pleiotropic effects - whereby genetic loci are associated with multiple traits - suggesting widespread connections between diseases at the molecular level (*2–5*). For example, cross-trait genetic associations have been reported for conditions where the affected organs differ but where there is some sharing of etiological mechanisms, such as the immune-mediated diseases (IMDs) (*6–9*). It has thus been recognised that the identification of pleiotropic associations could have the potential to help to define the genetic architecture of complex traits (*10*, *11*), provide insight into their evolutionary biology (*12*), and pave the way towards improved clinical care (*1*, *13*).

To date, however, it has not been possible to integrate and interrogate information from the full range of clinical phenotypes, as GWAS have focused on a relatively small number of traits and have often studied patients with only the most clear-cut diagnoses and uniform clinical manifestations. The availability of population-based cohorts, such as the UK Biobank (*9*, *14*, *15*), provides a unique opportunity to take a disease-agnostic perspective to investigate cross-trait genetic associations across a heterogeneous patient population. The UK Biobank has collected genetic and routine healthcare data from over 500,000 participants, including 531 diagnostic terms extracted from self-reported (SR) information, and 16,310 diagnostic terms from hospitalization episode statistics (HES). The latter are recorded using the tree of International Classification of Diseases, Tenth Revision (ICD-10) codes that categorizes diagnostic terms under 22 chapters corresponding to the major organ systems, such as “Diseases of the circulatory system”, or key disease classes, such as “Neoplasms”. Genome-wide single nucleotide polymorphism (SNP) information across all participants provides a powerful platform for identifying connections in genetic risk among common human diseases. However, there remain multiple analytical challenges in defining the structure of connections, including incomplete power, linkage disequilibrium (LD), underlying biological complexity, and a lack of resolution between diagnoses occurring as a direct cause of underlying pathology and those which may be secondary.

Here we address the challenges of working with phenome-wide data to resolve the genetic connectivity between disease traits in the UK Biobank. Making use of the hierarchical structure of standard disease classifications, we have characterised the overall genetic architecture of common disorders from routine healthcare data, assessing their genetic associations against prior GWAS and profiling their patterns of pleiotropy. We demonstrate that variants associated with multiple traits can be clustered based on the sets of phenotypes that they influence, revealing the presence of genetically determined, shared biological pathways that underpin different groups of diseases and that contribute differentially to shared genetic correlation between disorders. Moreover, these clusters can be used to re-define the relationship between medical conditions from a molecular perspective, thus providing insight of clinical value.

## Genome-wide associations with the UK Biobank routine healthcare data

To analyse genome-wide associations against all diagnostic terms in the UK Biobank SR and HES data sets in a simultaneous and hypothesis-free fashion, we employed our recent Bayesian analysis framework, TreeWAS, on 409,525 UK Biobank individuals with British Isles ancestry. This approach gains power for identifying phenome-wide associations by making use of the tree structure of routine healthcare data to estimate the evidence of association with at least one clinical diagnosis, summarised with the Tree Bayes factor statistic (BF_tree_), and to identify individual clinical phenotype codes that show association (*9*). Importantly, our approach allows for arbitrary distributions of effects over clinical codes. To enable comparison between variants, we simplify genetic effects into null, risk and protection for each code, integrating over a prior on effect size (see Methods). This results in strong correlation of BF_tree_ with the original implementation (Pearson rho = 1.00 and 0.99 in SR and HES, respectively; fig. S1). Of the 654,546 SNPs interrogated, we observed associations for 3.46% and 1.89% of them (log_10_ BF_tree_ ≥ 5) in the HES and SR data sets, respectively; and with 87.55% and 29.71% of the respective ontology terms showing evidence of association with at least one of these variants (PP ≥ 0.75) (Fig. 1A). Through permutation analysis (see supplementary materials) we estimated that at this threshold (log_10_ BF_tree_ = 5) we obtained a false positive rate of 5% (fig. S2). Evidence for association is highly correlated between the two data sets (Pearson *r* = 0.94; fig. S3), with SNPs rs9273363 and rs9272449 showing the strongest evidence of association in the HES (log_10_ BF_tree_ = 694) and the SR data sets (log_10_ BF_tree_ = 487), respectively. Both SNPs tag the *HLA-DRB1*03:01* allele (*r* = 0.62 and *r* = 0.73, respectively) in the major histocompatibility complex (MHC). The *HLA-DRB1*03:01* allele has been reported to have pleiotropic effects (*9*, *16*, *17*); we observe associations with 354 ICD-10 codes in the HES data set, including coeliac disease, type 1 diabetes, and systemic lupus erythematosus. After excluding the extended MHC (chr6:25,000,000-35,000,000), in total we identified 3,510 independent lead SNPs with a minor allele frequency (MAF) > 0.01 and LD trimmed (*r*^2^< 0.2) from the HES data.

**Figure 1.**
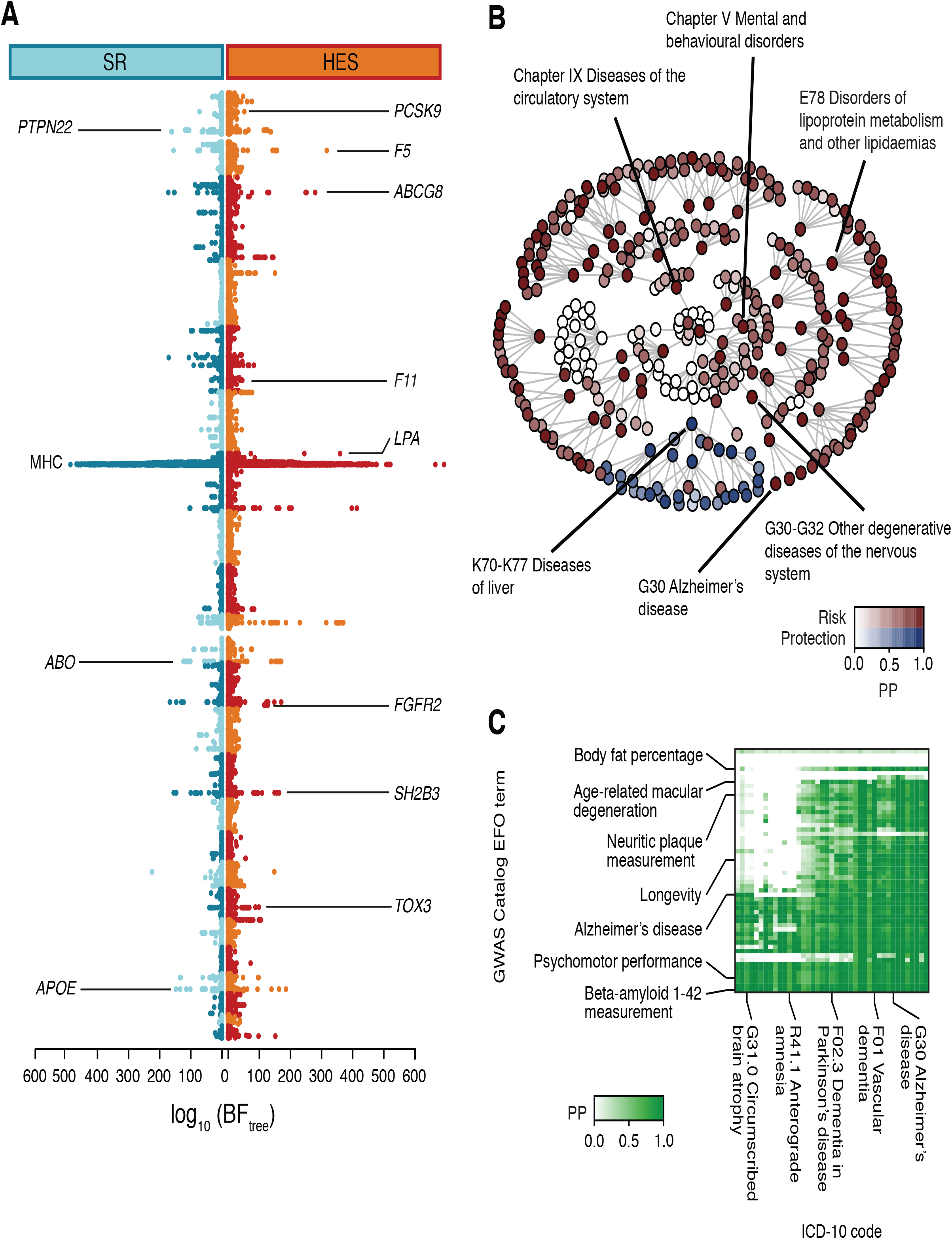
**(A)** Double Manhattan plot depicting evidence of association (log_10_ BF_tree_) across the SR (left) and HES (right) datasets. SNPs labelled with gene names exemplify notable associations to common human diseases and genetic evidence for therapeutic strategies to treat common diseases (see text). **(B)** Posterior decoding of genetic effect direction and strength of evidence for the rs4420638 SNP in the *APOE* locus as observed in the HES dataset ICD-10 classification tree. **(C)** Insert of the overlap, with a focus on Alzheimer’s disease, between previous GWAS (from NHGRI-EBI GWAS Catalog) and clinical associations in the UK Biobank HES data. Variants within the GWAS Catalog that are mapped to a specific Experimental Factor Ontology (EFO) term are tested, as a group, for evidence of association to the ICD-10 ontology. Colour indicates for each ICD-10 the posterior probability (PP) of association of the EFO variant set. Full analysis is available in the supplementary materials.

The extent to which even a single variant can impact a very broad range of biological and disease processes is demonstrated by the rs4420638 minor allele, which tags the *APOE*\**ε4* haplotype, the strongest genetic determinant for Alzheimer’s disease (*18*), and which is also associated with cardiovascular diseases (*19*) and lipid levels (*20*). We found the variant to confer risk for seven main clades within the ICD-10 ontology including those with parent nodes G30-G32 “Other degenerative diseases of the nervous system”; Chapter V “Mental and behavioural disorders”; Chapter IX “Diseases of the circulatory system”; E78 “Disorders of lipoprotein metabolism and other lipidaemias”; R41 “Other symptoms and signs involving cognitive function and awareness”; and Z95 “Presence of cardiac and vascular implants and grafts” (Fig. 1B, fig. S4). Unexpectedly, the same allele was also found to protect against one clade whose parent node is K70-K77 “Diseases of the liver” (Fig. 1B, fig. S4), demonstrating that implementing our approach across the HES data set can reveal previously unrecognised disease associations for even well-studied pleiotropic risk variants.

Patterns of genetic association across diseases can reveal unanticipated parallels and connections between genes with seemingly distinct functions. For example, GWAS have demonstrated that the nonsynonymous SNP rs3184504 (Trp262Arg) in *SH2B3* affects IMDs (*7*, *21*), but also cardiovascular traits (*4*, *5*, *19*) and cancer (*22*). We found associations with 207 ICD-10 codes for this SNP, including risk for multiple IMDs and cardiovascular disorders but protection against neoplasms such as C18 “Malignant neoplasm of colon” and C50 “Malignant neoplasm of breast” (fig. S5). Intriguingly, we found a similar pattern of associations for the rs2476601 (Arg620Trp) SNP in *PTPN22*, which has previously only been reported to confer risk for seropositive autoimmune diseases such as rheumatoid arthritis (*23*, *24*) and protection against Crohn’s disease (*25*). We observed risk associations between this SNP and 202 ICD-10 codes, the majority of which are IMDs; however, rs2476601 was also found to confer risk for cardiovascular disorders including I20 “Angina pectoris” and I21 “Acute myocardial infarction”, and protection against certain skin neoplasms (fig. S6). Both genes are expressed in haematopoietic cells, but whilst SH2B3 is an adaptor protein that affects growth factor and cytokine signalling (*26*), PTPN22 is a tyrosine phosphatase that regulates T and B cell receptor and toll-like receptor signalling (*27*). Nevertheless, specific perturbations of the *SH2B3* and *PTPN22* genes can have similar effects across multiple disorders, suggestive of an analogous impact on a shared biological process.

Other cross-trait association patterns reveal distinctions between genes thought to affect similar biological pathways. For example, for rs2289252 in the *F11* blood clotting factor locus, that is associated with venous thromboembolism (*28*), we observed a restricted set of diseases associations, only including I26.9 “Pulmonary embolism without mention of acute cor pulmonale”; I80.2 “Phlebitis and thrombophlebitis of other deep vessels of lower extremities”; Z86.7 “Personal history of diseases of the circulatory system”; and Z92.1 “Personal history of long-term (current) use of anticoagulants”.

However, whilst rs6025 (Arg534Gln), known as the Leiden mutation (*29*) which is found in the *F5* blood clotting factor gene, has also been reported to affect venous thromboembolism (*30*, *31*), we observed a much more diverse range of associations for this SNP. These include other vascular traits, such as I26-I28 “Pulmonary heart disease and diseases of pulmonary circulation” and I60-I69 “Cerebrovascular diseases”; infections (e.g. J14 “Pneumonia due to *Haemophilus influenzae*”); neoplasms (e.g. D17 “Benign lipomatous neoplasm”); paroxysmal neurological disorders (e.g. G43 “Migraine”); drug allergies (e.g. Z88.0 “Personal history of allergy to penicillin”); and surgical complications (e.g. T83 “Complications of genito-urinary prosthetic devices, implants and grafts”). Therefore, despite both SNPs influencing blood coagulation, their only partially overlapping disease association profiles suggest some disparity in the biological mechanisms they impact.

Further to associations with known pleiotropic loci, we could also detect the effect of genetic variants that have had implications for therapy, such as the low frequency rs11591147 SNP (Arg46His; MAF ≃ 2%) in the *PCSK9* locus that is correlated with reduced low-density lipoprotein cholesterol levels and coronary artery disease risk (*32*), and that has led to the efficacious trialling of PCSK9 blockers (*33*, *34*). We found a protective effect of this SNP against 8 SR diagnostic terms (log_10_ BF_tree_ = 72.21), and 67 ICD-10 codes (log_10_ BF_tree_ = 52.04), including E78.0 “Pure hypercholesterolaemia” and I25.1 “Atherosclerotic heart disease” (fig. S7). A protective effect of the minor allele of the rs11209026 SNP (Arg381Gln) in *IL23R* was also observed, affecting 21 ICD-10 codes (log_10_ BF_tree_ = 19.24), which included inflammatory bowel disease-related clinical nodes (K50-K52 “Noninfective enteritis and colitis”) and a weak evidence of association (PP = 0.73) in psoriasis (ICD-10 code L40 and child nodes including L40.5 for psoriatic arthritis). This allele has been previously reported to protect against multiple IMDs and correlates with reduced IL-23-mediated signalling (*6*, *7*, *25*, *35*). Notably, IL-23 blockade has shown promise in the treatment of Crohn’s disease, psoriasis and psoriatic arthritis (*36–38*), consistent with the observed ICD-10 code associations. Thus, characterising the extent and nature of pleiotropic genetic effects could help to reveal targets amenable to drug repositioning strategies, by providing a rationale for which diseases could be treated with the same therapeutics, as well as which could not.

We next sought to assess the overlap of our results with previous GWAS. The NHGRI-EBI GWAS Catalog (*39*) has compiled over 15 years of GWAS results and mapped diseases and trait terms to the Experimental Factor Ontology (EFO). Among the 41,445 SNPs present in the GWAS Catalog and genotyped or imputed in the UK Biobank cohort, we found evidence for association (log_10_ BF_tree_ > 0) with TreeWAS for 48.4% and strong evidence for association (log_10_ BF_tree_ ≥ 5) for 8.2%. GWAS Catalog variants are enriched among SNPs with increasing levels of evidence of association (fig. S8; odds ratio (OR) of 2.91 for SNPs with log_10_ BF_tree_ ≥ 5), and depleted of SNPs with a weak level of association (log_10_ BF_tree_ between 0 and 1; OR = 0.70, *P* = 2.86 x 10^-218^).

These results confirm that the UK Biobank has substantial power to detect genetic associations across a wide range of clinical traits. However, one of the major challenges in comparing results between the GWAS Catalog and the UK Biobank is the lack of standard for mapping between ontologies (here EFO and ICD-10). This problem can be at least partially solved by identifying factors in the two ontologies linked through genetic association. To establish such a mapping, we calculated, for each set of GWAS Catalog SNPs mapped to a specific EFO term, their joint evidence of association across the HES data (see Methods). We found that 41.73% of the EFO terms had evidence of association (log_10_ BF_tree_ > 0). We then used posterior decoding to identify where in the ICD-10 classification tree the set of SNPs for each EFO term had an effect (Fig. 1C, fig. S9). Often, we found evidence for multiple linkages between ontologies. For example, the SNP set associated with the Alzheimer’s disease EFO term in the GWAS Catalog was associated, within the UK Biobank, with the HES G30 “Alzheimer’s disease” code (PP = 1.0; Fig. 1C), as well as with 26 other ICD-10 codes including G30 child nodes and numerous other mental disorder diagnostic terms (F00-F09 “Organic, including symptomatic, mental disorders” and its child nodes). Conversely, G30 was also found to be associated (PP ≥ 0.99) with SNP sets affecting EFO terms for apolipoprotein E isoform E2, beta-amyloid 1-42, imaging and psychomotor measurements, demonstrating the potential for genetics to define meaningful connections between ontologies for classifying diseases, biological terms and biomarkers.

## The structure of genetic pleiotropy in the UK Biobank HES data

To quantify the structure of genetic pleiotropy in the UK Biobank, we determined the relationship between the evidence of association for the 3,510 lead SNPs and the number of ICD-10 codes associated with it (PP ≥ 0.75). We find that 97.92% of associated SNPs affect more than one diagnostic term, with the common SNP (MAF ≥ 5%) with the most associations being rs13107325 (Ala391Thr/Ser) in *SLC39A8*, affecting risk for 303 terms across 9 Chapters of the ICD-10 ontology, consistent with previous observations for this SNP being associated with multiple phenotypes (*3*, *39*). Overall, variants with greater evidence of association affect a larger number of diagnostic terms (*ρ* = 0.21, *P* < 1.0×10^-16^, Fig. 2A). However, we also observed variants with very strong evidence of association (log_10_ BF_tree_ > 20) that affect only a small number of phenotypes. For example, rs2981575 and rs4784227 (both log_10_ BF_tree_ > 90) localise (on different chromosomes) near *FGFR2* and *TOX3,* respectively, and are associated with the same 19 nodes in the ICD-10 ontology, all related to breast cancer (including C50 “Malignant neoplasm of breast” and its child nodes) and procedures such as Z51.1 “Chemotherapy session for neoplasm”. These SNPs have a similar association profile, displaying a strong evidence of association with a high precision in the phenotypes affected, and this likely reflects a strong similarity in the biological pathways they influence. Overall, we found that 66.81% of the 3,510 lead variants were associated with the top node of the ICD-10 classification tree and 68.58% of the SNPs were associated with at least 2 of the 22 Chapters of the ICD-10, providing evidence that most genetic variants affecting risk to a diagnostic term will often also affect risk to other terms distant in the ontology.

**Figure 2.**
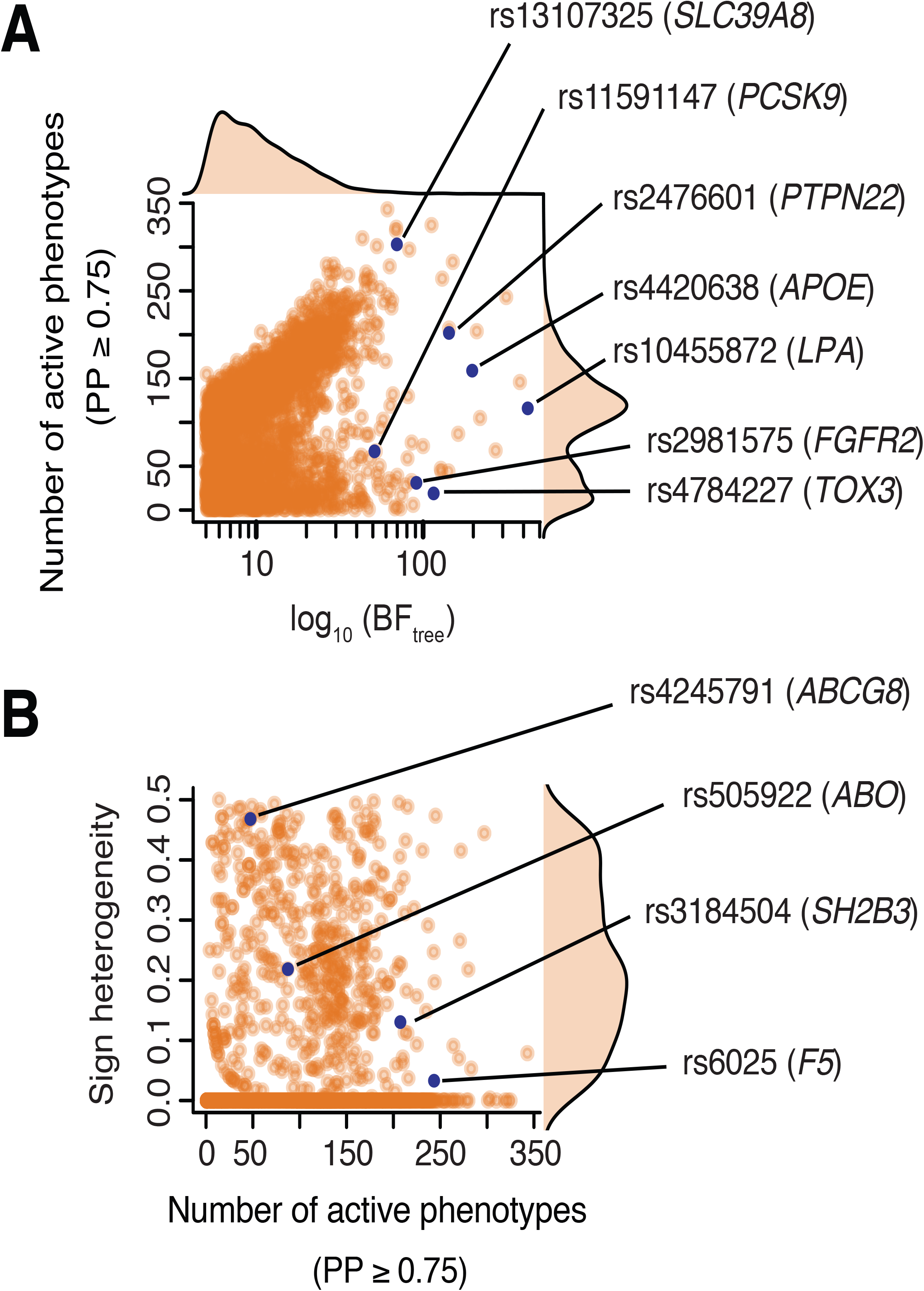
Genetic pleiotropy across common human diseases in the HES data set. **(A)** Relationship between the evidence of association of a SNP in the disease ontology (BF_tree_) with the number of phenotypes associated with the SNP with non-zero effect sizes (PP ≥ 0.75). **(B)** Sign heterogeneity in SNP effects across associated phenotypes. Sign heterogeneity per SNP is defined as the minimum between the ratio of phenotypes where the minor allele is risk and the ratio of phenotypes where the minor allele is protective.

We next considered the extent to which variants show a combination of risk and protection across different classes of disorder. For each associated SNP we calculated sign heterogeneity, defined as the minimum of the ratio of phenotypes for which the minor allele confers risk (PP ≥ 0.75) and the ratio of phenotypes for which this allele promotes protection (range 0 to 0.5). We found that 85.38% of associated SNPs show no sign heterogeneity, whilst for the 14.62% that do, the degree of sign heterogeneity was approximately uniformly distributed (Fig. 2B). For example, the rs4245791 SNP in *ABCG8* was found to be associated with 47 nodes (log_10_ BF_tree_ = 127.69) and had a sign heterogeneity of 0.47, with the minor allele protecting against disorders of the gallbladder (22 ICD-10 codes) and increasing risk to hypercholesterolaemia and cardiovascular diseases (25 ICD-10 codes). SNPs that reduce risk for some diseases but increase it for others may provide more challenging therapeutic targets for which, however, putative side effects of the therapeutic intervention could be predicted.

## Decoding cross-trait associations through hierarchical SNP clustering

Across independently associated variants we observed several repeated patterns of risk and protection suggestive of distinct genes modulating the same underlying biological processes. To test this hypothesis formally, we calculated, for every pair of variants, a Bayes factor comparing a model in which they share the same profile, to a model in which they are independent (see Methods). We then used hierarchical clustering to define groups of variants with similar profiles (see Methods, Fig. 3A, fig. S10) and for each cluster we computed a joint posterior decoding to identify associated diagnostic terms. After filtering for variants with strong evidence in the HES data, we identified 629 distinct clusters with sizes ranging from 1-51 SNPs. Overall, 50% of SNPs occurred in the largest 161 clusters of 7 or more SNPs each. For example, the previously mentioned highly pleiotropic rs3184504 and rs2476601 SNPs in *SH2B3* and *PTPN22*, respectively, both lie in clusters of size one, while the less pleiotropic *PCSK9* rs11591147 SNP lies in a cluster of 19 variants. The diagnostic code with the greatest number of distinct clusters showing association is N19.8 “Chronic renal failure”, followed by I50.0 “Congestive heart failure”, and the cluster with the greatest number of SNPs affects 33 nodes.

**Figure 3.**
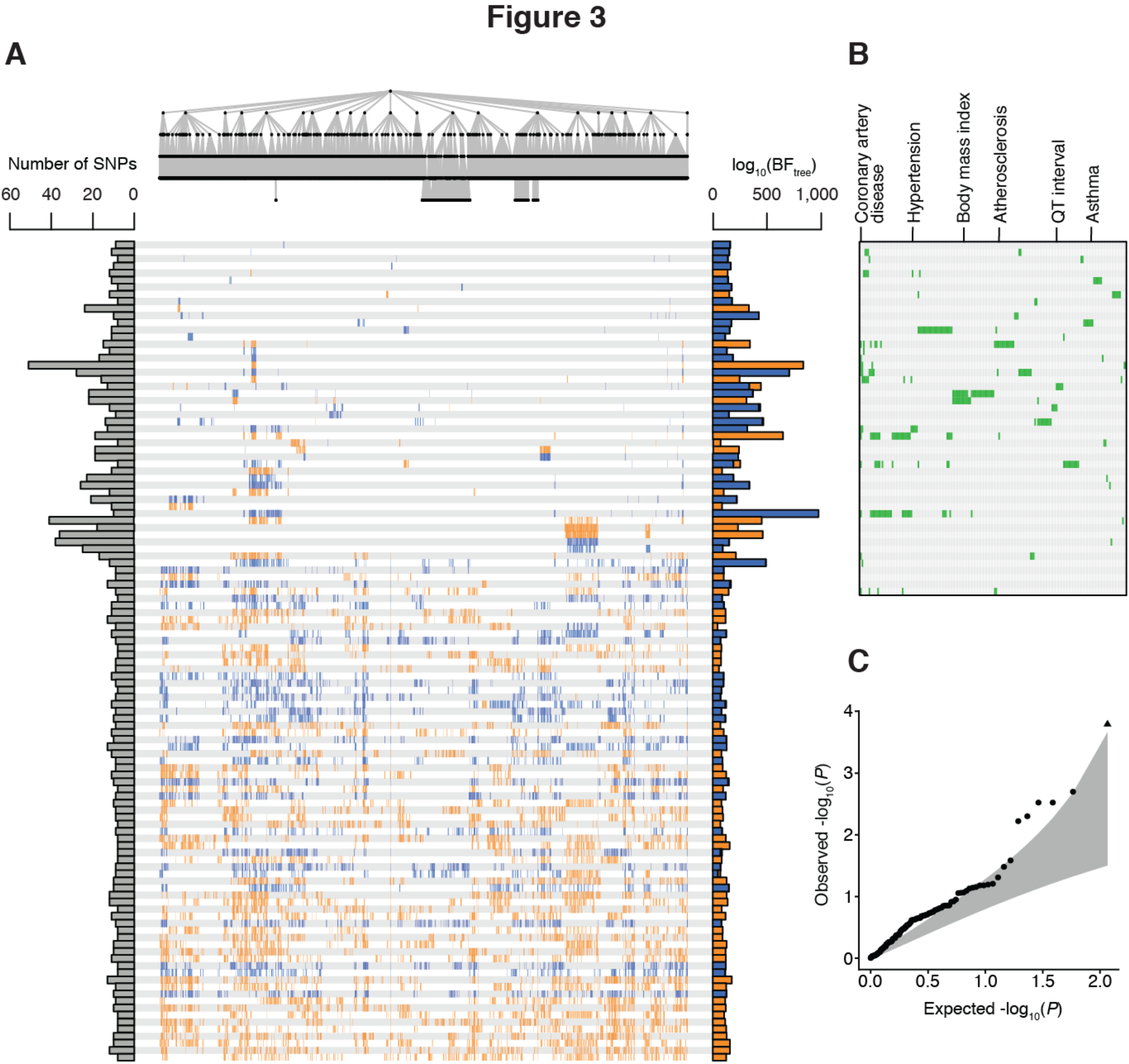
Genetic risk profiles across common diseases in the HES data set. **(A)** Posterior decoding for 116 risk profiles with at least 8 SNPs identified from clustering of 3,510 variants. The tree on the top depicts the ICD-10 ontology where each node represents an ICD-10 code. Below the tree each row represents a risk profile, where bars depict associations (PP ≥ 0.75) to ICD-10 codes, and colours relate to the direction of the effect (orange is risk; blue is protection). The barplot on the left shows the number of SNPs present in each cluster and the barplot on the right the estimated BF_tree_ for the cluster with colour-coding based of proportion of associations that are risk (orange) and protective (blue). **(B)** Extent to which SNPs present in each cluster are enriched in GWAS Catalog phenotypes. Results are shown for the top 66 clusters as the remaining ones do not show evidence of enrichment, with the exception of clusters 93, 103 and 115 with GWAS Catalog terms ‘cognitive decline measurement’, ‘schizophrenia’ and ‘bone density’, respectively. Each column represents one GWAS Catalog EFO term. **(C)** Enrichment of GO biological process terms in cluster SNP sets. For each cluster SNP set we calculate enrichment statistics for all GO terms and record the minimal *P*-value observed across all terms. We then, for each cluster, calculate an empirical *P*-value which is the proportion of times the minimal GO term *P*-value is smaller than those observed by randomly generating SNP sets from background of the same size (see Methods).

Each cluster represents a potentially distinct biological mechanism or pathway conferring risk for common diseases, with distinct patterns of potential co-morbidity. To investigate this hypothesis, we assessed enrichment of variants within each cluster among SNPs reported previously in the GWAS Catalog (at the level of EFO terms) and to gene ontology (GO) terms for biological processes, by mapping individual SNPs to genes in a 100 Kb window around each SNP (see Methods). We find 139 clusters that show overlap with EFO terms (permutation *P* < 0.05; Fig. 3B) and, 69 clusters with evidence for enrichment in GO terms (permutation *P* < 0.05; Fig. 3C). For example, Cluster 12, containing 8 SNPs, associated with ICD-10 codes in the blocks J40-J47 “Chronic lower respiratory diseases” and J30-J39 “Other diseases of upper respiratory tract”, as well as branches containing the clinical diagnosis for asthma and nasal polyps (ICD-10 codes J45 and J33, respectively). The SNPs in this cluster were enriched for GWAS Catalog SNPs reported for the EFO terms asthma (Fig. 3B), eosinophil percentage of granulocytes, eosinophil percentage of leukocyte, eosinophil count, neutrophil percentage of granulocytes (permutation *P* < 0.05) and GO terms related to interleukin-5 production (*P* = 9.7×10^−8^). Genes in the vicinity of these SNPs that are linked to the significant GO terms include *GATA3*, *IL1RL1* and *IL33*. Given, that GATA3 inhibition is an effective treatment for asthma (*40*), genes affected by other SNPs in this cluster could serve as potential targets for the same biological pathway.

A cluster-based approach to dissecting genetic risk can reveal the multiple different processes and pathways that contribute to any single clinical endpoint. To illustrate this, we considered the single most common code within the UK Biobank HES data, I10 “Essential (primary) hypertension” (for which there are 24.37% of individuals with at least one record of this code). We observed 69 distinct clusters with strong association to the code (PP ≥ 0.75), which each affect between one and 698 ICD-10 codes (PP ≥ 0.75). Among these clusters, one affects hypertension only; 19 are strongly associated with type 2 diabetes and obesity; 14 are associated with hypercholesterolaemia, angina and myocardial infarction; 23 are associated with chronic kidney disease; four are associated with disorders of the gallbladder and bile duct such as cholangitis, and 30 are extremely broadly-acting, affecting over 300 ICD-10 codes, including mental health, tobacco use and alcohol abuse (Fig. 4). This diverse range of different clinical phenotypes associated with each of these many clusters suggests that multiple, heterogeneous biological mechanisms can underpin the development of any single common, complex disease. Clusters affecting hypertension are typically small (median 5 SNPs), and we see no significant GO enrichment for any, indicating the need for variant-level (as opposed to gene level) annotation of molecular function and biological process.

**Figure 4.**
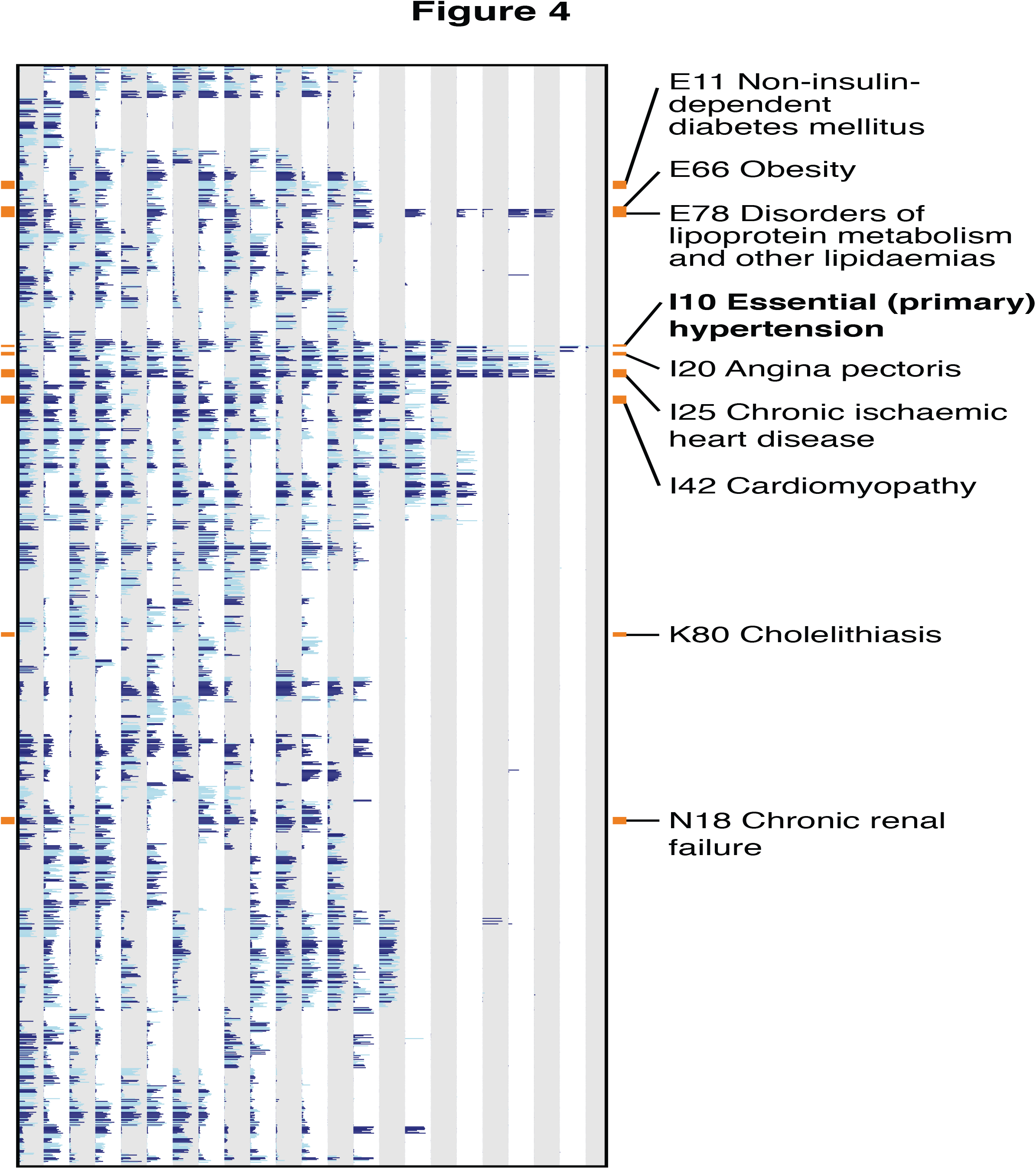
Genetic risk profiles associated with hypertension. 23 risk profiles for clusters with at least 5 SNPs associated with the ICD-10 term I10 “Essential (primary) hypertension” (PP ≥ 0.75). For each cluster (columns), the relative posterior across 1,206 codes (where at least one of the clusters shows evidence for association with PP ≥ 0.75 and where annotations are observed) is shown. Light and dark blue indicate ICD-10 codes at the category level. Terms mentioned in the text are highlighted.

## The genetic ontology of common human diseases

Finally, we considered the extent to which patterns of genetic risk are consistent with the ICD-10 ontology or could be well represented by any other single hierarchical ontology. For example, IMDs are widely distributed across the ICD-10 ontology, not least because they affect various tissues and organs across the body, and genetic risk scores for these diseases are highly precise in identifying the focal disorder within the UK Biobank (*9*). However, IMDs are also well known to share many genetic risk factors (*6–8*), indicating overlapping underlying biological risk processes, consistent with this we observed that 265/629 clusters are associated with risk for at least two distinct IMDS (PP ≥ 0.75, see Methods).

We first measured the evidence, for each cluster of SNPs, that they impact more than a single clade (a given node and all descendant nodes) within the ICD-10 ontology, by calculating a Bayes factor comparing models with single and arbitrary numbers of clades of activity. Among the 629 clusters, 94.43% show compelling evidence (log_10_ BF ≥ 5) for multiple clades (Fig. 5A), with the estimated number of switches in active state along the topology ranging from 2 to 736. However, we also found overwhelming evidence that the ICD-10 ontology describes the correlation structure of association better than a star-ontology (log_10_ BF > 1,000). These results indicate that the ICD-10 ontology provides a partial, but incomplete, match to the structure of genetic risk.

**Figure 5.**
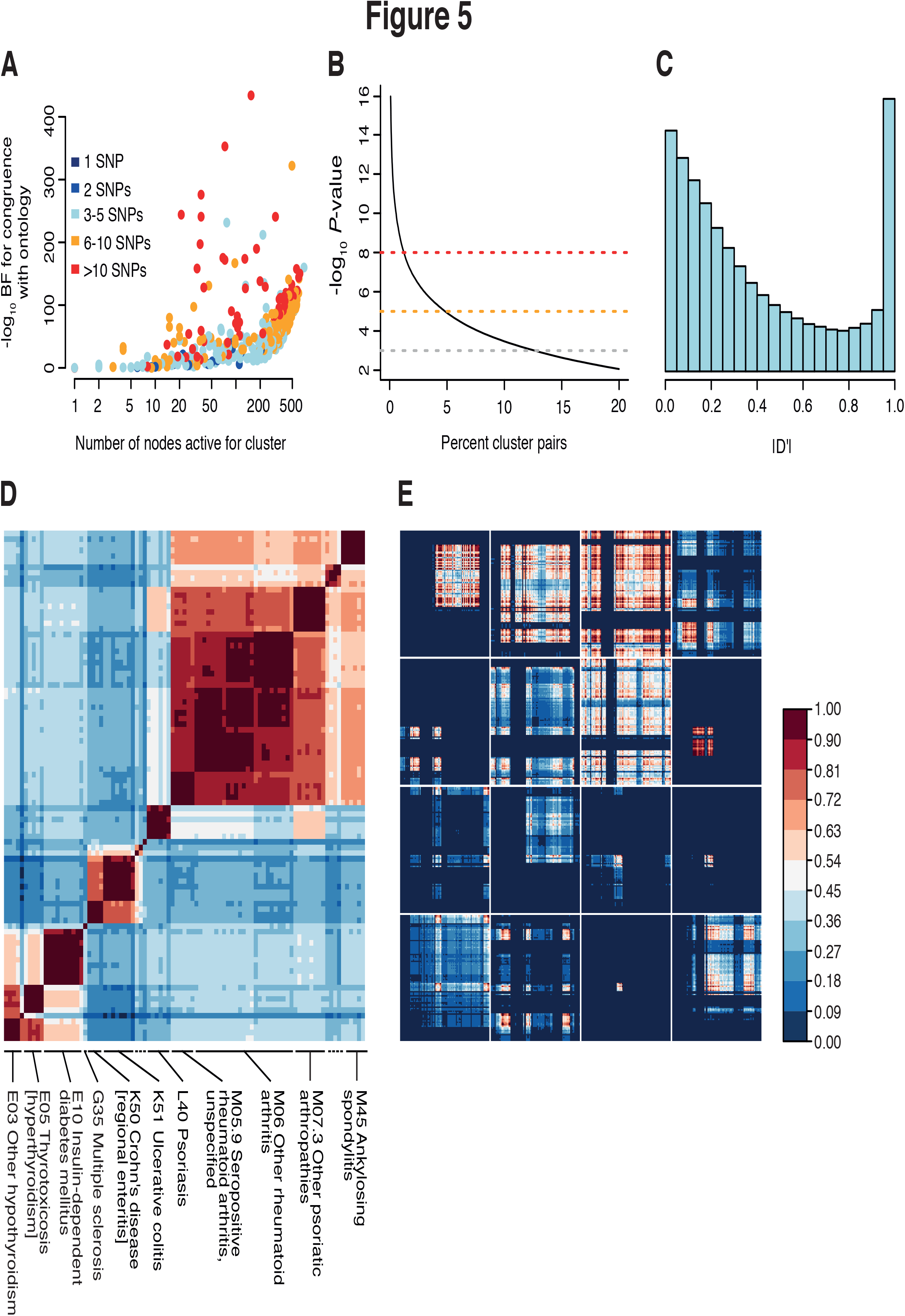
Genetics and disease ontology. **(A)** The relationship between the breadth of phenotypic activity for clusters, number of SNPs in a cluster, and evidence (log_10_ BF) for the cluster affecting > 1 clade in the ICD-10 ontology. **(B)** Histogram of the estimated phenotypic disequilibrium, |D’|, between pairs of clusters. **(C)** The percentage of pairs of clusters showing a given level of evidence (*P*-value from LRT assuming Chi-squared distribution with one degree of freedom) for patterns of association that are inconsistent with any hierarchical ontology. Coloured lines indicate *P* < 10^-3^ (grey), *P* < 10^-5^ (orange) and *P* < 10^-8^ (red). **(D)** Heatmap showing the overlap in posterior profiles across clusters (weighted by variants in cluster) for 91 clinical codes relating to common IMDs. **(E)** Genetic covariance for 16 clusters that each affect at least one of the ICD-10 code L40 “Psoriasis” child nodes (PP ≥ 0.75).

We next sought to ask whether any single hierarchical ontology could adequately capture the structure of association. For every pair of clusters, we calculated the evidence for departure from hierarchical consistency, by comparing models in which the effects of the variants across clinical codes are consistent with a tree (only three out of the four possible combinations of active/inactive at the two clusters are present) to one in which they can have arbitrary patterns (see Methods). In doing so, we estimate a quantity we refer to as ‘phenotypic disequilibrium’ between clusters, which is analogous to the |D’| statistic measure of linkage disequilibrium (*41*). We find that 12.6% of pairs of clusters show strong evidence (LRT, *P* < 0.001, see Methods) for the presence of all four combinations of activity across phenotypes, which is incompatible with any single hierarchical ontology (Fig. 5B). The distribution of phenotypic disequilibrium is skewed towards low values, with only 15.7% of pairs of clusters having |D’| ≥ 0.9 (Fig. 5C). We therefore conclude that patterns of genetic risk across common diseases typically exhibit combinations of risk and/or protection that are inconsistent with any single hierarchical ontology.

Although our results falsify the notion of a single hierarchical ontology of underlying genetically determined pathophysiological processes, nevertheless, they can be used to decompose the well-established genetic covariance between traits, such as for IMDs (*6–8*). For example, among 91 ICD-10 codes corresponding to IMDs we observe patterns of overlap in genetic risk consistent with previous findings (*6–9*, *42*) (Fig. 5D). However, the covariance can be decomposed into the contributions of individual clusters, which have diverse structures. Notably, we find 41 clusters associated with child nodes of L40 “Psoriasis” (PP ≥ 0.75), which show a wide variety of patterns of covariance (Fig. 5E), consistent with the observed heterogeneity in symptoms, comorbidities and response to treatment among patients (*43*, *44*).

## Discussion

Our approach has enabled a unified interrogation of the structure of genetic risk for multiple disease traits. We have demonstrated how the architecture of genetic risk for complex disorders, such as hypertension and IMDs, consists of distinct groups of loci that impact specific sets of disorders. These results indicate the presence of underlying biological pathways, whose dysregulation can affect risk for a variety of disorders and which, through the identification of the underlying genes and molecular mechanisms, offer multiple routes to therapeutic intervention. For example, the presence of 69 distinct SNP clusters associated with hypertension suggests that there may be as many different genetically determined pathways driving this condition that are potentially amenable to drug targeting. However, prioritisation or exclusion of clusters for further investigation in a therapeutic context would likely depend on their full profile of cross-trait associations. For instance, we find evidence that for multiple clusters, typically those comprising a small number of variants, there is a widespread and often complex pattern of risk. This suggests that therapeutic perturbation of the biological pathways corresponding to these clusters would result in a lack of specificity and potential off-target or adverse effects, whereas the subset of clusters with more restricted or uniform association profiles would likely reveal more promising drug targets.

The overall architecture of genetic risk that we observed mirrors previous findings relating to the structure of pathophysiological pathways implicated in common diseases, such as the presence of a small number of genes acting as master regulators of risk with a widespread impact across multiple traits, and many specialised genes with more disease-specific effects (*3*, *10*, *11*). Unexpectedly, we found that few loci show evidence for sign heterogeneity, even among loci affecting a large numbers of disorders, though there are notable exceptions, such as variants in *APOE* and the ABO and Colton red blood cell antigen systems. Theoretically, both a trade-off between early-benefit and later-disease risk or mutation-selection-drift balance can maintain genetic variance for complex diseases (*45*). Here, we find little evidence for trade-offs at the level of common disease, though early life benefits may be manifest through processes other than avoiding risk for disease (such as through sexual selection) or from challenges (such as infection or starvation) that are rare in western society. Rather, our results are more compatible with complex disease arising from dysregulation of underlying quantitative traits, where small genetic perturbations can either be beneficial or deleterious depending on their genetic background (*45*). Future fine-mapping of causal variants and inference of their ancestral state will indicate whether new mutations always increase the risk of disease or, as would be expected from a stable quantitative trait, are balanced in terms of effect direction.

From a clinical perspective, the identification of genetic pathways of risk raises the potential for defining a biologically meaningful substructure to the diagnosis, prognosis and treatment of common disorders. An individual’s risk for any single disease spans multiple dimensions, each one with a specific set of potential comorbidities (*46*). Individuals who reach disease may therefore possess differential dysregulation across underlying dimensions, manifest as different patterns of comorbidity and potentially differential responsiveness to treatment options. The identification of genetic clusters with similar association profiles, and the elucidation of the molecular mechanisms they correspond to, thus suggest the possibility for discovering dimension-specific biomarkers and therapeutic targets. However, an important implication of our findings is that the phenotypic consequences of different dimensions do not typically form a simple hierarchical ontology. Rather, genetics reveal a complex connectivity between diseases that is reflected through multiple different combinations of conditions.

Finally, we note that our approach has highlighted key analytical challenges, including the need to handle covariates and to model non-genetic associations between traits adequately. Appropriately modelling the impact of genetics on the longitudinal nature of disease and the complex interplay between factors, whilst also coping with the heterogeneity that arises from variable data recording and clinical practices, remains an important and unsolved problem. Moreover, ultimately, each locus has a unique biology, and hence our approach of identifying variant clusters is essentially a statement of ignorance: the full integration and interpretation of clinical, quantitative and molecular phenotypes will require new approaches for data sharing, aggregation and analysis.

## ACKNOWLEDGEMENTS

This research has been conducted using the UK Biobank Resource (application number 10625). This work uses data provided by patients and collected by the NHS as part of their care and support.

## Funding

This research has been conducted with the support of the Wellcome Trust (100956/Z/13/Z and 090532/Z/09/Z to G.M. and 100308/Z/12/Z to L.F.), the Danish National Research Foundation (L.F.), the Medical Research Council (MC_UU_12010/3 to L.F.), the Oak Foundation (OCAY-15-520 to L.F.), the National Institute for Health Research (NIHR) Oxford Biomedical Research Centre (BRC) to L.F., the Wellcome Trust/Royal Society (204290/Z/16/Z to C.A.D.) and the Li Ka Shing Foundation (to G.M.).

## Competing interests

G.M. is a cofounder of, holder of shares in, and consultant to Genomics PLC, and is a partner in Peptide Groove LLP. Peptide Groove has licensed HLA typing technology to Affymetrix Ltd. The other authors declare no competing financial interests.

## Author contributions

A.C. and G.M. performed the analyses with contributions from C.A.D. and L.F. L.F. and G.M. conceived the study. A.C., C.A.D., L.F. and G.M. wrote the manuscript.

## Data availability

The data shown in this paper are available as part of the supplementary materials.

